# Dietary Lipid Oxidization Products Alter Growth, Adiposity and Gut Microbial Ecology in Prepubertal Porcine Model

**DOI:** 10.1101/2022.03.06.483183

**Authors:** Folagbayi K. Arowolo, Kent A. Willis, Ibrahim Karabayir, Oguz Akbiligic, Morgan Blaser, Jeffrey Booth, Joseph F. Pierre, Dhanansayan Shanmuganayagam

**Affiliations:** Biomedical & Genomic Research Group, University of Wisconsin-Madison, Madison, WI; Molecular and Environmental Toxicology Center, University of Wisconsin-Madison, Madison, WI; Department of Pediatrics, Division of Neonatology, College of Medicine, The University of Alabama at Birmingham, Birmingham, AL; Department of Health Informatics and Data Science, Parkinson School of Health Sciences and Public Health, Loyola University Chicago, Maywood, IL; Department of Econometrics, Kirklareli University, Kirklareli, Turkey; Department of Pediatrics, College of Medicine, University of Tennessee Health Science Center, Memphis, TN; Agricultural Research Stations, University of Wisconsin–Madison, Arlington, WI; Department of Animal and Dairy Sciences, University of Wisconsin–Madison, Madison, WI; Department of Surgery, University of Wisconsin School of Medicine and Public Health, Madison, WI

**Author notes:** **Corresponding Author:** Dr. Dhanansayan Shanmuganayagam, Ph.D., Department of Surgery, Department of Animal and Dairy Sciences, University of Wisconsin-Madison, 1675 Observatory Drive, Rm 666, Madison, WI 53706, E-mail: –.

**Keywords:** Swine, High-Fat Diet, Lipids, Gastrointestinal Microbiome, Adiposity

## Abstract

Elevated levels of dietary fats in westernized diets, associated with increased risk of obesity and other chronic diseases, are increasingly consumed by children in the United States. Cooking practices such as high heat frying and increased use of oxidizable sources of fats have introduced high levels of lipid oxidation products (LOPs) into these diets. The effects of these highly reactive dietary compounds on human biology are largely unstudied, especially in the gut where these compounds are likely present at higher concentrations. Given that the gut microbiome can be influenced by dietary components and then in turn have a systemic impact, we investigated the effects of consuming LOPs on gut bacterial and fungal communities and on growth and body composition during the prepubertal period in a porcine model. The presence of LOPs in the high fat diet reduced growth and body fat gain in the model. The gut microbiome was uniquely altered by both high fat and the presence of LOPs, with notable changes in the abundances of *Turicibacterales, Spriochaetales, RF39, Lactobacillales and Erysipelotrichales*. The mycobiome was dominated by *Kazachstania*, a porcine specific yeast, which was only minimally influenced by the dietary regimen. Application of machine learning identified dietary fat and LOPs as strong predictors of body fat. The genus *Methanobrevibacter* was the key microbial predictor of body fat. This study highlights the need for further studies on the biological effects of LOPs which have become ubiquitous in human, livestock and pet diets in developed countries.

## 1. INTRODUCTION

Over the past half-century, increased consumption of the Western diet – composed of lipid-rich and highly-processed foods – has correlated with increased incidence of chronic diseases.^1–3^ While consumption of fats in general increased over this period, a dramatic change in the types of fats consumed also occurred. Intake of saturated fats (e.g., animal fats: lard, butter and tallow) declined and then stabilized,^4^ while intake of oils and fats rich in polyunsaturated fatty acids (PUFAs; e.g., vegetable oils: corn, soybean and canola oil) has continued to rise.^5^ For example, in North America, while animal fat intake declined by 36% between 1963 and 2003, vegetable oil intake increased by 136%, and is projected to be 218% higher by 2025 when compared to intake in 1963. This public trend was driven by the dogma that saturated fats are primarily responsible for increased risk of chronic diseases,^6,7^ while PUFAs are healthier alternatives, or even protective.^8,9^ However, the latter assumption has complicated implications.

PUFAs are highly susceptible to oxidation due to the presence of unsaturated bonds, and easily form reactive lipid oxidation products (LOPs). Decades of research have shown endogenously formed LOPs to be modulators of immune responses^10^ and mechanistic drivers of chronic diseases,^11,12^ especially atherosclerosis.^13^ However, what has been largely ignored in the human health field is that the increased use of PUFAs, along with increased food processing and changes in cooking methods,^14^ have introduced increasing quantities of LOPs into the Western diet,^15^ increasing the influx of exogenous LOPs into the body. Fats and oils rich in PUFAs, especially when subjected to high temperatures such as those used in deep frying,^14^ rapidly oxidize to form primary LOPs such as lipid hydroperoxides, that in turn can further undergo conversion into a wide spectrum of secondary and tertiary LOPs, including aldehydes, ketones, and other polymeric compounds.^16^ With a third of Americans daily frequenting fast food establishments,^17–19^ known for their broad use of deep frying, the consumption of food products rich in LOPs continues to rise.^20^ This trend has especially swept over to children.^21,22^ The fraction of meals children consume at fast food restaurants increased by 296% between 1977 and 1996.^23^ The proportion of PUFAs consumed by children also increased by 73% in the same period.^24^ Given that several chronic diseases such as atherosclerosis initiate in childhood^25^ and risk factors measured in childhood and adolescence have been shown to be better predictors of the severity of atherosclerosis than risk factors measured in young adults,^26^ it is important to understand the impact of consuming LOPs during childhood development.

Given the recent scientific interest in the link between gut microbiome and adiposity as it relates to risk of chronic diseases,^27^ in this study, we utilize a porcine model due to its translational advantages^28,29^ to study the effects of consuming LOPs in prepubertal period on the gut bacteriome, mycobiome and body composition. While a number of studies have examined the biological effects of high-fat diets on the gut microbiome and body composition,^30–33^ no study thus far has given any attention to the oxidative status of fats being consumed. Here, we also utilize machine learning to develop a predictive model to explore the association between specific changes in gut microbiome and body composition in the pig model.

## 2. MATERIALS AND METHODS

### 2.1 Analysis of Lipid Oxidation Products in Study Oils

Corn oil (Columbus Vegetable Oils, Des Moines, Iowa, USA) was used to provide fat content for the high fat diet, but without significant addition of LOPs, while spent restaurant oil (i.e., used for cooking) from local restaurants (Sanimax Industries Inc, DeForest, WI, USA) was used to provide fat content for the high fat diet containing high amounts of LOPs. The concentration of primary LOPs, specifically lipid peroxides, in the fat sources were measured using a modified version of an iron-based spectrophotometric method as previously described,^34,35^ and reported as peroxide value (PV). Secondary LOPs, such as aldehydes and ketones, in the fat sources were measured using p-anisidine value (AV) test as previously described.^36^ The quantities of select LOPs were determined in the study oils using a liquid chromatography-mass spectrometry (LC-MS) method.^37^

### 2.2 Animal Study Design

The experiments involving animals were conducted under protocols approved by the University of Wisconsin-Madison Institutional Animal Care and Use Committee in accordance with published National Institutes of Health and United States Department of Agriculture guidelines. Twenty-four domestic piglets bred and maintained at the University of Wisconsin Swine Research and Teaching Center [40,000 sq ft, specific pathogen-free (SPF) facility] were weaned at 3 weeks of age (initial body weight: 8.2 ± 0.2 kg) were block-randomized and allocated to three different dietary regimens at 1 month of age until ~5.6 months of age (**Figure 1**): (1) **Low fat diet** (LF, 9% calories from fat), (2) **High fat diet with negligible/low LOPs** (HF, 44% calories from fat), (3) **High fat with high LOPs diet** (HF+LOPs, 44% calories from fat. The piglets were assigned to and group-housed in pens (4 pens per diet regimen), ensuring equal distribution of sex (1 female:1 male) and matching average body weight (at weaning) across all pens. The piglets were placed on a grain-based acclimation diet for a week before the assigned dietary regimens were initiated. Pigs were given *ad libitum* access to feed and water; feed intake was measured and monitored throughout the study. The experimental diets were formulated and prepared by University of Wisconsin Arlington Research Station Feed Mill to meet nutritional requirements for the various dietary phases of a developing pig (**Supplementary Table 1**).^38^ The feed was prepared every two weeks and stored at 4°C to minimize changes in the oxidation status of diets over the duration of the experimental feeding period. The pigs self-adjusted their daily feed-intake to accommodate for the calorie-density of the dietary regimen such that daily calorie intake did not differ between the dietary groups (**Supplementary Figure 1**).

**Figure 1.**
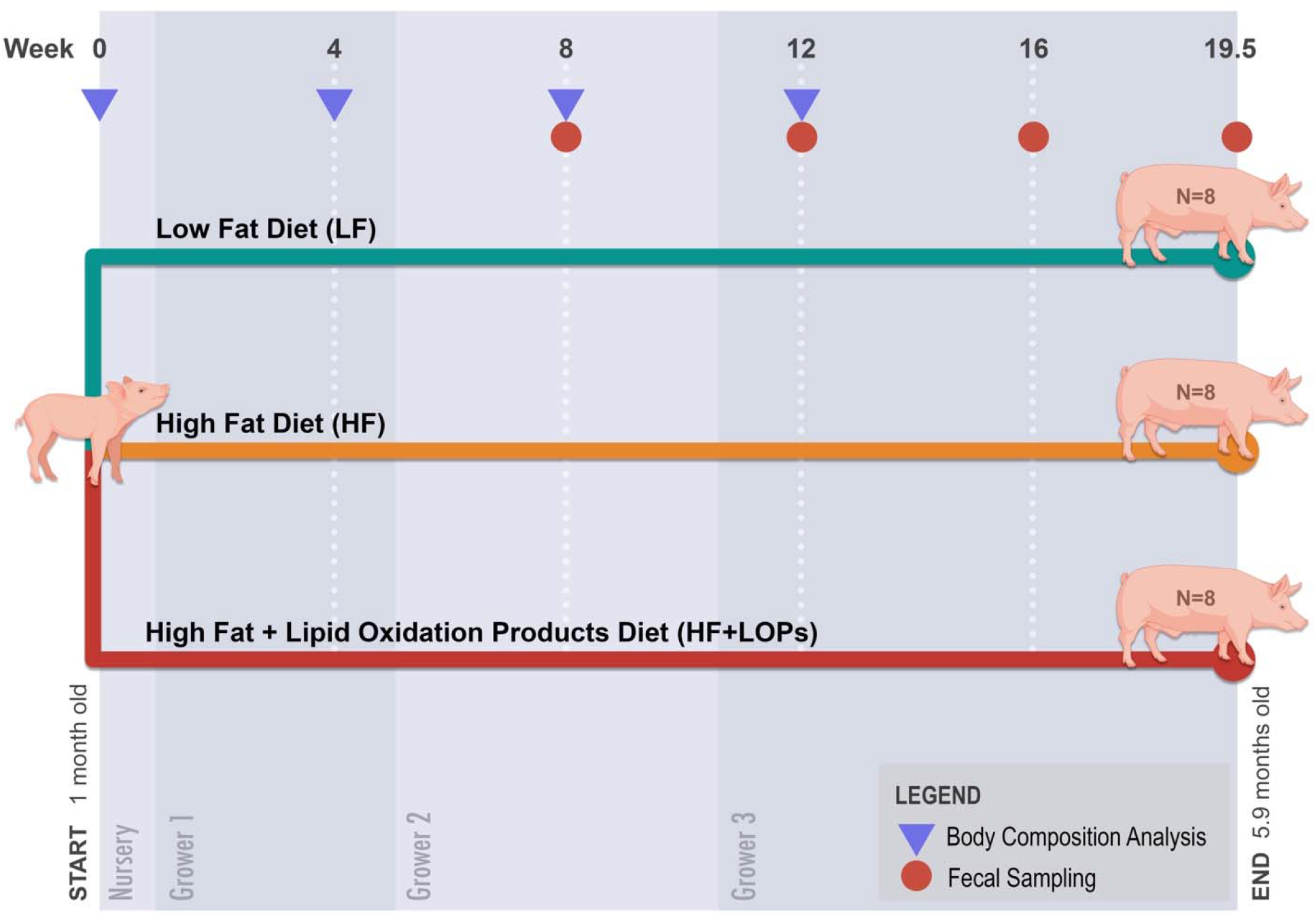
Study Design. Twenty-four domestic pigs were assigned to three diet regiments at 1 month of age and maintained until end of study at 5.9 months of age (approximate age at which pigs reach sexual maturity at our breeding facility). While the relative proportionality of nutrients and parameters of interest of the (i.e., dietary fat (LF=9% of calories from fat, HF = 44% of calories from fat) and lipid oxidation products(LOPs)) remained consistent throughout the study across dietary groups, the feed formulations were changed at specific age points according to standard practices to meet nutritional requirements of a developing pig (diet formulation details are provided in Supplementary Table 1); these nutritional phases are denoted by differences in background shading and labelled “Nursery,” “Grower 1,” “Grower 2,” and “Grower 3.” Body weights and feed intake were recorded weekly and bi-weekly, respectively. Symbols denote body composition analysis (blue triangles) and fecal samples (red circles) available for use in study.

Body weights were measured weekly for the entire duration of the study. Body composition of the pigs was determined monthly by dual energy x-ray absorptiometry (DXA), until the size of the pigs became prohibitive. On days of the DXA scans, fecal samples were also obtained from freshly voided fecal matter (up to 5g) or rectally using a sterile cotton swab when samples were difficult to obtain. Fecal samples were stored in a conical tube at −80°C until DNA extraction. It should be noted that although attempts to collect fecal samples prior to the start (at timepoint 0) and one month after start of study diet regimen were made, the samples could not be used for microbiome analysis due to collection challenges.

### 2.3 Body Composition Analysis

Body composition was determined using DXA (GE Lunar Prodigy, GE Healthcare, Madison, WI). Prior to the DXA imaging, the pigs were fasted for 24 hours (access to drinking water was not limited). Anesthesia was induced (Narkomed 2, Drager, Telford, PA) with 5–7% sevoflurane or isoflurane and maintained with 2–4% isoflurane or sevoflurane delivered through a nosecone to prevent pig movement during DXA scan (10 to 20 min). Anesthetic depth was confirmed by absence of ocular reflex, loss of muscle tone, absence of pedal reflex, and deep breathing. Pulse-oximeter was used to monitor heart rate and blood oxygen saturation. Then, the pigs were placed in a sternal recumbency position and imaged using the appropriate scan mode (Small Animal [5–20 kg], Adult Standard [20–46 kg] or Adult Thick [above 46 kg]), according to the body weight of each pig. Postscan analysis provided estimates of % total body fat (TBF, %), truncal body fat (TrBF, %), extremity body fat (EBF, %), total bone mass (TBM, %), truncal bone mass (TrBM, %), extremity bone mass (EBM,%), total lean mass (TLM, %), truncal lean mass (TrLM, %), and extremity lean mass (ELM, %).

### 2.4 DNA extraction and Illumina MiSeq sequencing

Porcine fecal samples were resuspended in 500 mL of TNES buffer containing 200 units of lyticase and 100 mL of 0.1/0.5 (50/50 volume) zirconia beads and incubated for 20 minutes at 37°C. Following mechanical disruption using ultra-high-speed bead beating, 20 mg of proteinase K was added to all samples and incubated with agitation overnight at 55°C. Total DNA was extracted using the chloroform isoamyl alcohol, and total DNA concentration (per mg stool) was determined by qRT-PCR. Purified DNA samples were sent to the Argonne National Laboratory (Lemont, IL) for amplicon sequencing using the NextGen Illumina MiSeq platform, utilizing 16S rRNA MiSeq for bacteria and archaea and parallel ITS2 rDNA sequencing for fungi. Blank samples passed through the entire collection, extraction and amplification process remained free of DNA amplification.

### 2.5 Bioinformatics

Sequencing data were processed using QIIME 1.9.1.^39^ Sequences were first demultiplexed, then denoised and clustered into operational taxonomic units (OTUs). For bacteria, sequences were aligned via PyNAST ^40^ and taxonomy was assigned using the RDP Classifier.^41^ For fungi, sequences were aligned, and taxonomy was assigned using the UNITE (dynamic setting) database.^42^ Processed data were then imported into Calypso 8.84^43^ for further data analysis and visualization. On import, all sequences aligning to chloroplasts or cyanobacteria were discarded. For bacteria, any samples with less than 3,000 sequence reads were discarded, resulting in the removal of two samples from downstream analysis. For fungi, samples with less than 300 sequence reads were discarded, resulting in the removal of 6 samples from downstream analysis. A read depth of 300 has been previously validated as sufficient for capturing the diversity of the porcine mycobiome.^44^ Principal components analysis (PCA), principal coordinates analysis (PCoA) and redundancy analysis (RDA) plots of Bray-Curtis dissimilarity distances were utilized to visualize multivariate beta diversity.^45^ Statistical significance of beta diversity clustering was then assessed using RDA and permutational multivariate analysis of variance (PERMANOVA) followed by permutational analysis of multivariate dispersions (PERMDISP2) to assess the homogeneity of group variance (distance to centroid). To assess alpha diversity, bacterial samples were first rarified to a depth of 9,683 reads and fungal samples were rarified to a depth of 2,988 reads then the Shannon and Chao1 diversity indices were calculated and differences assessed using analysis of variance (ANOVA). Univariate analyses using analysis of comparison of microbiomes (ANCOM), ANOVA, core microbiome analysis and mixed effect regression, with sex as a random effect, were performed to quantify the differences in the relative abundance of specific microbial taxa.^46^

### 2.6 Statistical Analysis

Student’s t-test was used to determine statistical significance of differences in primary and secondary LOPs between fat sources utilized in the diets; data are reported as means ± standard deviation (SD). To test the significant effects of diet regimen on body weight and body composition, a fixed effects ANOVA model was fit using the PROC MIXED function of SAS software (version 9.4, SAS Institute Inc., Cary, NC, USA).^47^ The model included terms to define fixed effects (diet regimen and sex), random effects (housing pen), and repeated measures (subject). Correlations between measurements collected from the same animal over time were modeled using an autoregressive model. The models were fit using untransformed data and the resulting residuals were used to determine if ANOVA assumptions of constant variance and normality were sufficiently met. Significance was determined using Type III tests and was accepted at *p* < 0.05. Data are reported as least-square means (LSM) ± standard error of the mean (SEM).

### 2.7 Machine Learning Based Modeling

For machine learning models, a Light Gradient Boosting Machine (LightGBM)^48^ was used to explain the variation of the three dependent variables (TBF, EF, TrBF) as a non-linear function of sex, diet type and determined OTUs, which came from the level of family and genus, were present in at least 2 samples in the dataset. LightGBM is a relatively novel yet very powerful ensemble algorithm that can be used to carry out both classification and prediction (regression). It is based on building weak decision tree learners and iteratively adding more learners to minimize the prediction error in the previous weak learner.^48^ Due to limited data size, the effect of outliers in dataset may have great influence on the predictive models. Therefore, in addition to the original dataset, experiments were separately run on both with and without outlier detection for comparison. Ultimately, three main datasets with sample sizes of 22, 20 and 19 were utilized. The use of two hierarchical ranks of OTUs for the analyses and three dependent variables led to the generation of six different analyses for each main dataset. To detect possible overfitting with machine learning modeling, we implemented a cross validation procedure and reported only test results. In order to be consistent with training phase between folds, and due to limited sample size, the Leave One Out Cross Validation (LOOCV) method was implemented. Bayesian optimization algorithm was utilized for parameter tuning of the LightGBM regression. In the model building phase, early stopping rule was implemented to avoid overfitting. Mean squared error (MSE) with 90% confidence interval was calculated using the predicted values after LOOCV **(Table 2)**. For clarity, true values and predictive values were plotted. Lastly, variable importance analysis was performed to identify the effective variables on building models.

## 3. RESULTS

### 3.1 Analysis of Lipid Oxidation Products in Study Oils

PV in corn oil and spent restaurant oil were 1.62 ± 0.11 mmol O_2_/kg (3.24 ± 0.21 mEq O_2_/kg) and 14.34 ± 1.15 mmol O_2_/kg (28.68 ± 2.31 mEq O_2_/kg), respectively, while AV in corn oil and spent restaurant oil were 3.35 ± 2.24 and 45.13 ± 3.77, respectively. That is, primary (PV) and secondary (AV) LOPs were 9-fold (*p* < 0.001) and 14-fold (*p* < 0.001) greater, respectively, in spent restaurant oil than in corn oil. Analysis of secondary LOPs, specifically C9-C11 aldehydes, in the study oils, showed that only three out of the fourteen LOPs analyzed were detectable in corn oil and in low concentrations, while all of the LOPs analyzed were present in spent restaurant oil and in high concentrations (**Table 1**).

**Table 1.** Concentration of Secondary Lipid Oxidation Products, Specifically C9-C11 Aldehydes, in Study Oils.

|  | **Corn Oil**<br>**(mg/L)** | **Spent Restaurant Oil**<br>**(mg/L)** |
| --- | --- | --- |
| Hexanal | 1.8 | 176.9 |
| Acrolein | ND | 175.1 |
| 2-Hexenal | ND | 53.5 |
| 2,4-Decadienal | ND | 47.5 |
| Pentanal | 0.8 | 42.4 |
| 2-Heptenal | 0.3 | 39.9 |
| 4-hydroxynonenal<br>(4-HNE) | 2.0 | 15.0 |
| Nonanal | ND | 14.4 |
| 2-Octenal | ND | 10.1 |
| 2,4-Heptadienal | 0.1 | 10.1 |
| 2-Nonenal | ND | 6.2 |
| Octanal | ND | 5.4 |
| 2-Pentenal | ND | 3.2 |
| 2,4-Nonadienal | ND | 1.2 |

ND = Not detectable

### 3.1 Body Weight and Body Composition

Significant differences in body weight gain (*p* < 0.0001) were observed in the pigs under the dietary regimens over the course of the pre-pubertal study period. By week 13, pigs fed the high-fat diet (HF) had a higher (12.6%,*p* < 0.0012) body weight than those fed the low-fat diet (LF) (**Figure 2**). During this period, the weight of pigs fed the high fat diet containing lipid oxidation products (HF+LOPs) was not significantly different from those of LF and HF pigs. By week 15, the body weight of pigs fed HF and HF+LOPs were higher (13.3%,*p* < 0.0001 and 7.1%, *p* < 0.02, respectively) than LF pigs. This trend continued to the end of the study; the body weight of HF pigs was 13.1% (*p* < 0.0001) higher than LF pigs. However, the body weight of HF+LOPs pigs diverged and was 4.9% lower (*p* < 0.02) than those of their HF counterparts at the end. It should be noted that, as expected, sex of the pigs had a significant effect (*p*<0.001) on their body weight during development, but there was no significant interaction between sex and diet (data not shown).

**Figure 2.**
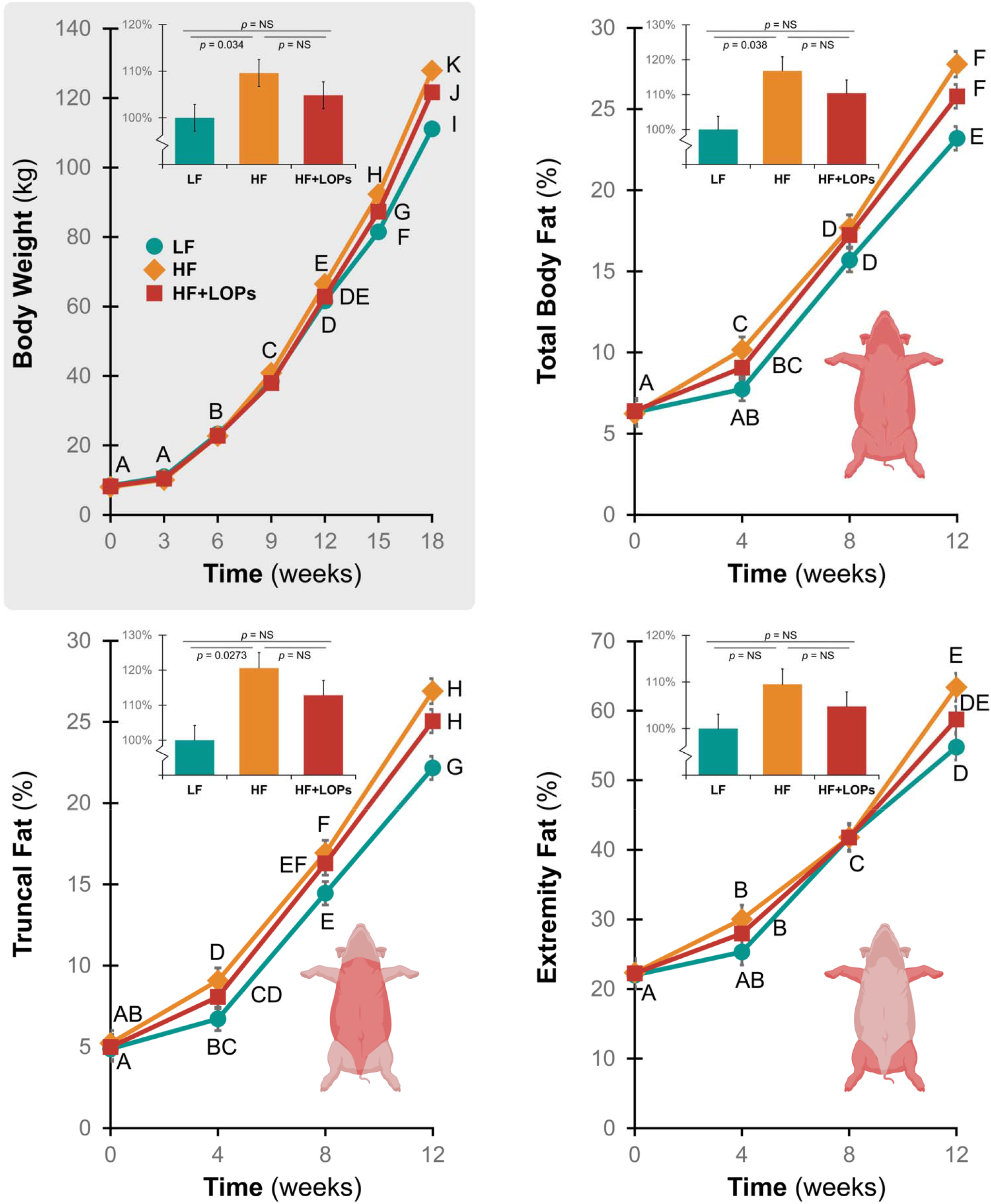
Effects of Dietary Lipid Oxidation Products on Body Weight and Adiposity (Total and Regional) in the Prepubertal Period. The line graphs show changes in total body weight and adiposity (total, truncal and extremity) over the course of the study. Data is presented as least square means ± SEM. Means without any common letter differ, *p* < 0.05. Some error bars, due to their relatively small magnitude, are hidden by the symbols used to denote the parameter in the graphs. Adiposity, determined by dual energy x-ray absorptiometry (DXA), could only be determined up to week 12, after which point the size of the pigs (particularly the chest girth) prevented the use of the medical imaging platform. Adiposity is expressed as the percentage of fat mass compared to the total mass of the region of interest. The inset bar graphs show the overall effect of the diet regimens (least square means ± SEM) on the parameters, expressed relative to the effects in LF pigs. While HF produced significantly higher changes in body weight and adiposity (except in extremity fat) in the pigs, the presence of LOPs consistently blunted this effect.

Significant differences in total and regional body composition (adipose, bone and lean mass) were observed in the pigs under the dietary regimens during the pre-pubertal study period (**Figures 2 and 3**). At study completion, pigs fed HF had a 19.7% higher TBF (*p*<0.01) and 21.4% higher TrBF (*p*<0.01), when compared to those fed LF. There were no significant effects of diet regimen on EBF (**Figures 2**). Pigs fed HF also showed significantly higher % bone mass (TBM and TrBM, but not EBM, **Figures 3**). The presence of LOPs seemed to have a slightly more pronounced increase in % bone mass, including EBM. However, these effects when compared to those of HF were not statistically significant (**Figures 3**). Overall, only HF pigs showed a significant difference in lean mass (TLM and TrLM, but not ELM) when compared to LF pigs.

**Figure 3.**
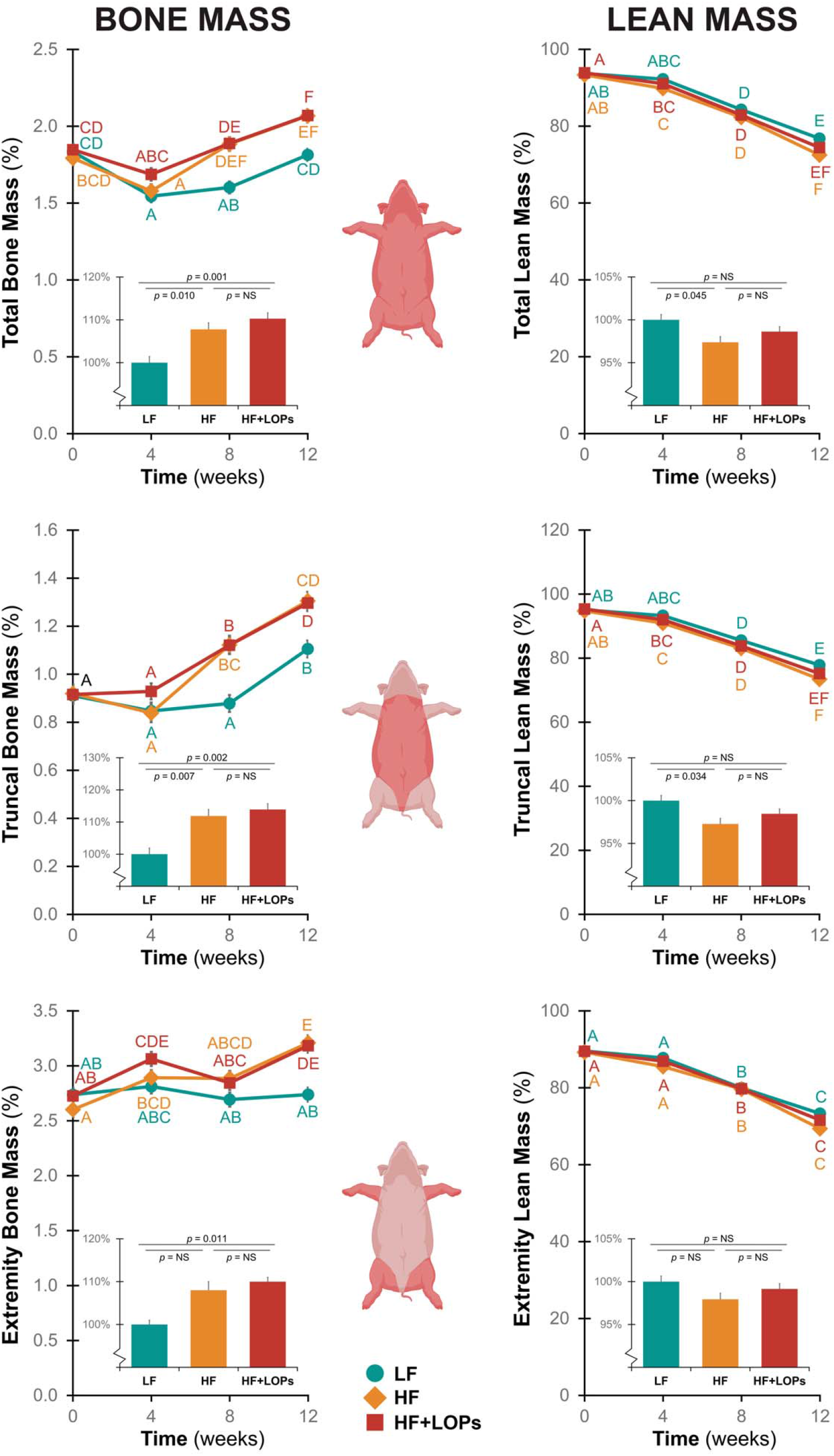
Effects of Dietary Lipid Oxidation Products on Bone and Lean Mass in the Prepubertal Period. The line graphs show changes in bone and lean mass (total, truncal and extremity) over the course of the study. Data is presented as least square means ± SEM. Means without any common letter differ, *p* < 0.05. Some error bars, due to their relatively small magnitude, are hidden by the symbols used to denote the parameter in the graphs. The bone and lean mass are expressed as a percentage to the total mass of the region of interest. The inset bar graphs show the overall effect of the diet regimens (least square means ± SEM) on the parameters, expressed relative to the effects in LF pigs. In general HF and HF+LOPs produced significantly higher increases in bone mass as the pigs developed. HF, and not HF+ LOPs produced significantly lower changes in total lean mass, most likely attributable to effects on lean mass in the truncal region.

### 3.2 Gut Bacteriome Taxonomy and Diversity

To assess temporal changes in bacterial community dynamics, we utilized samples collected at two-months (early timepoint) and five months (late timepoint) after the start of the dietary regimens. Two months of diet regimen resulted in distinct difference in bacteriomes between each of the dietary group (**Figure 4**). However, by five months on the diet, these differences were less significant, and multivariate analysis demonstrated that the beta diversity (inter-sample variance) of the gut bacteriomes more closely converged (Bray-Curtis dissimilarity, PERMANOVA, *R^2^* = 0.357, *p* = 0.00033, PERMDISP2, *p* = 0.0161; RDA, variance = 62.68, *f*= 1.91, *p* = 0.001, **Figure 4a**). Community convergence was in part driven by decreased alpha diversity (intrasample variance) observed over the study period as the animals aged (Shannon, ANOVA *f* = 7.5, *p* = 6.4 x 10^-5^ and Chao1, ANOVA *f* = 4, *p* = 0.0057, **Figure 4b**). To gain a more granular understanding of differences in microbial composition, we first visualized differences in relative abundance of bacterial orders using ANOVA adjusted for false discovery rate (FDR, **Figure 4c**), which highlighted differences in relative abundance between the orders *Turicibacterales, Spriochaetales, RF39, Lactobacillales* and *Erysipelotrichales*. We then utilized more rigorous univariate techniques to quantify differences in relative abundance of bacterial genera. Broadly, mixed effect regression identified *Clostridiaceae species* (spp.), *Turicibacter*, *Streptococcus, Megasphaera, Prevotella, Ruminococcaceae spp., Treponema, RF39 spp., Lachnospiraceae spp., Clostridiales spp., Lactobacillus, Clostridium* and *S247 spp.* as the most significantly altered genera (all FDR-adjusted *p* < 1×10^-14^, **Supplement Table 2**). Most of the ten significantly altered genera identified by ANCOM uniformly decreased in abundance over the course of the experiment, most notably *Bacteroides, Bulleidia, Faecalibacterium, Lachnospira, Streptococcus* and *Turicibacter* in each diet group (**Supplement Table 3**). However, the abundance of *Epulopiscium* remained unchanged in animals exposed to the LF diet and was nearly undetectable in animals receiving LOPs. *Barnesiellaceae spp*. exhibited the opposite effect and remained at higher abundance in animals receiving LOPs. We identified bacterial genera with biomarker potential utilizing LEfSe (**Supplement Figure 2**).

**Figure 4.**
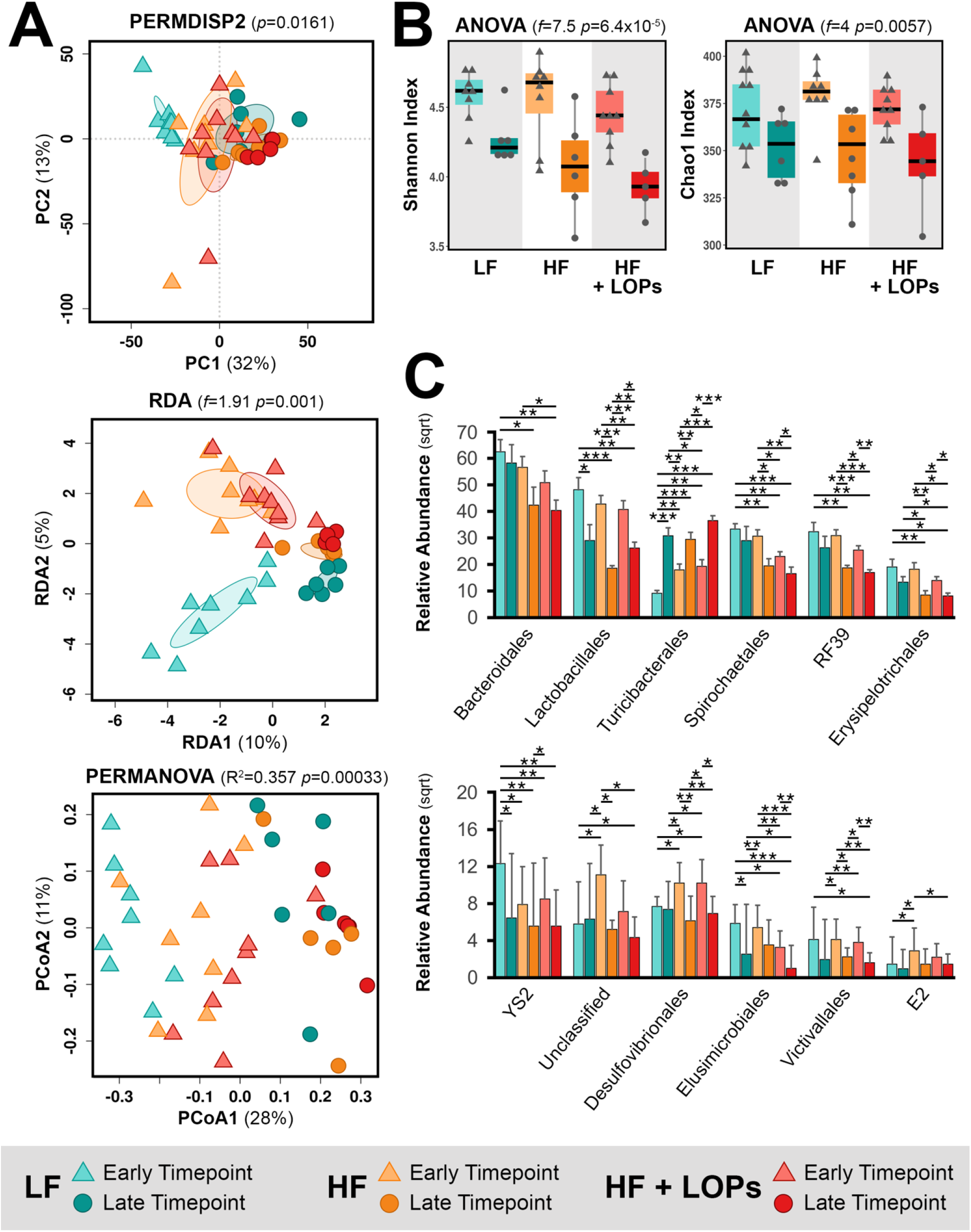
The bacterial microbiome converges with time on all three diets. **a** Multivariate analysis demonstrate that the bacterial microbiome converges with time [Principal components analysis (PCA), permutational analysis of multivariate dispersions (PERMDISP2) *p* = 0.0161; redundancy analysis (RDA), variance = 62.68, *f*= 1.91, *p* = 0.001; principal coordinates analysis (PCoA), permutational multivariate analysis of variance (PERMANOVA) *R^2^* = 0.357, *p* = 0.00033)]. **b** Diversity decreases with time (Shannon, ANOVA *f* = 7.5, *p* = 6.4 x10^-5^ and Chao1, ANOVA *f* = 4, *p* = 0.0057). **c** Relative abundance of bacterial orders alter with time (ANOVA false-discovery rate adjusted \**p* < 0.05, \*\**p* < 0.01, \*\*\**p* < 0.001).

At the end of the experiment, multivariate analyses demonstrated that bacterial communities, while convergent over the course of the experiment, remain distinct (Bray-Curtis dissimilarity, PERMANOVA, *R^2^* = 0.239, *p* = 0.004, PERMDISP2, *p* = 0.0497, RDA, variance = 16.36, *f* = 1.17, *p* = 0.005; **Figure 5a**). Community diversity, however, converged (Shannon, ANOVA, *f* = 7.5, *p* = 0.12 and Chao1, ANOVA, *f* = 0.18, *p* = 0.83, **Figure 5b**). Core microbiome analysis identified a core microbiome of 70 genera, with 4 unique genera in LF fed pigs and 2 unique genera, *Acinetobacter* and *Rheinheimera*, in pigs exposed to dietary LOPs (**Figure 5c**). Similarly, ANOVA identified *Paraprevotella, Methanobrevibacter* and *Bulledia* decreased in LOPs-fed pigs with a corresponding bloom in *Enterobacteraceae* spp. (**Figure 5d**). Mixed effect regression identified *Methanobrevibacter*, S247 spp., *Treponema,* Clostridiaceae spp., RF39 spp., *Ruminococcus* and *Prevotella* as the most significantly differentially abundant genera (all FDR-adjusted *p* < 0.0001, **Supplement Table 4**). More precisely, ANCOM identified *Epulopiscium* as the only significantly altered genera, which was decreased in animals exposed to LOPs (**Supplement Table 5**).

**Figure 5.**
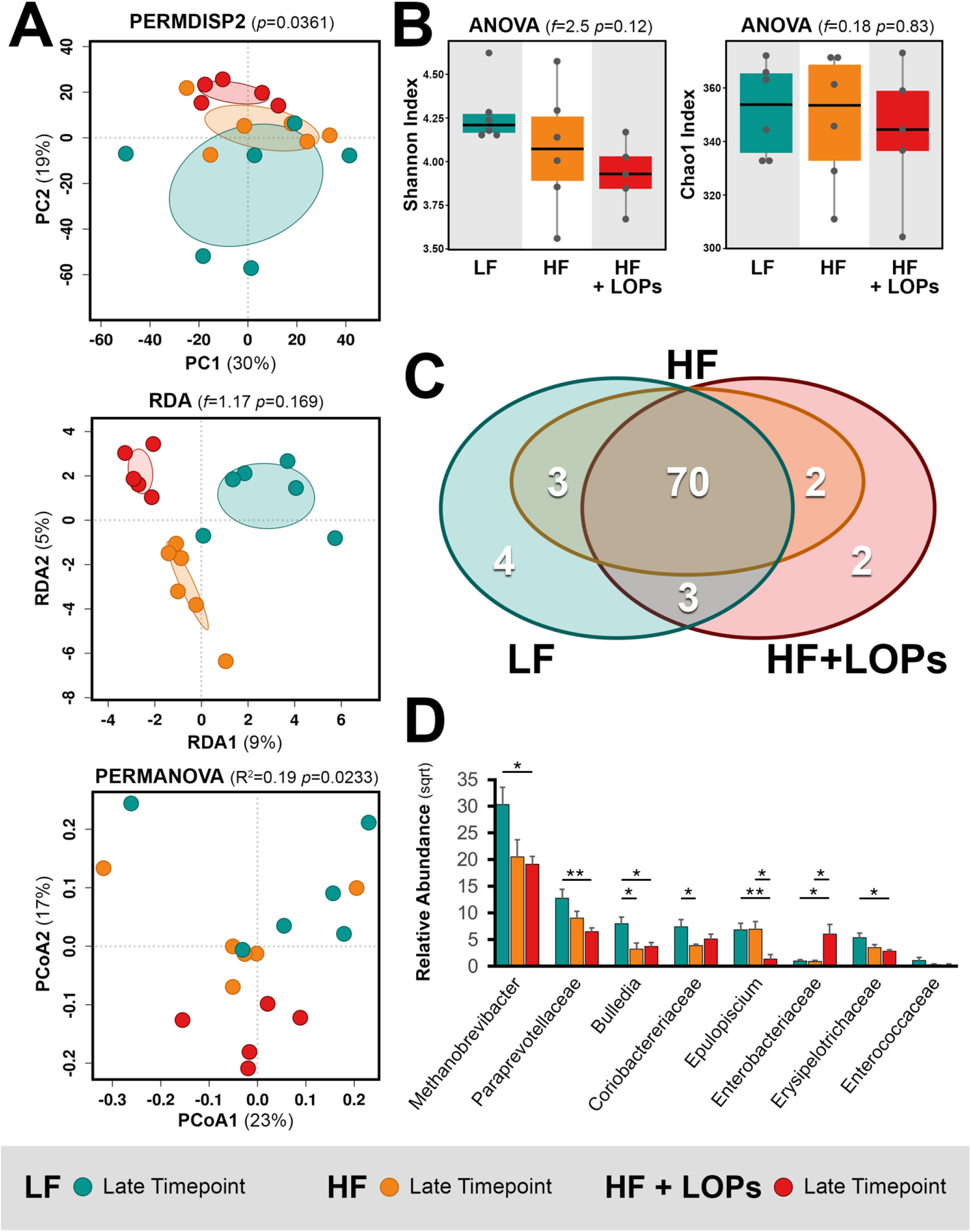
Lipid oxidation product in the diet alters microbial ecology. **a** Lipid oxidation products produce a porcine stool microbiome that diverges from animals on high fat diet [Principal components analysis (PCA) of Bray-Curtis dissimilarity, permutational analysis of multivariate dispersions (PERMDISP2) *p* = 0.0361; redundancy analysis (RDA), variance = 62.68, *f* = 1.17, *p* = 0.169; principal coordinates analysis of Bray-Curtis dissimilarity, permutational multivariate analysis of variance (PERMANOVA) *R^2^* = 0.19, *p* = 0.0233)]. **b** Bacterial diversity is not significantly altered (Shannon, ANOVA *f* = 7.5, *p* = 0.12 and Chao1, ANOVA *f*= 0.18, *p* = 0.83). **c** Core microbiome analysis shows unique more genera with lipid oxidation exposure. **d** Relative abundance of fungal genera (ANOVA false-discovery adjusted \**p* <0.05, \*\**p*<0.01, \*\*\**p* <0.001).

### 3.3 Gut Mycobiome Taxonomy and Diversity

Fungal gut communities in mammals are sparse in comparison to the more abundant and diverse bacteriome.^49,50^ As previously described,^51^ the porcine mycobiome in our study was also dominated by a single taxa, *Kazachstania*, which is a yeast genus in the family Saccharomycetaceae (**Figure 6**). Variability in community structure then is driven by minor taxa that account for only a limited percentage of sequence reads. Similar to observations in bacterial communities, two months of diet regimen resulted in distinct fungal community composition that converged over the course of the experiment. Multivariate analyses demonstrated that fungal communities in LF-fed pigs diverged from communities in HF-fed pigs (Bray-Curtis dissimilarity, PERMANOVA, *R^2^* = 0.239, *p* = 0.004, PERMDISP2, *p* = 0.0497, RDA, variance = 16.36, *f* = 1.17, *p* = 0.005). While the diversity of fungal communities in LF-fed pigs expanded as assessed by Shannon diversity, the diversity of communities in HF-fed pigs contracted dramatically (ANOVA, *f* = 6, *p* = 0.0046, **Figure 6a**), but not as assessed by Chao1 (ANOVA *f* = 1.2, *p* = 0.33). ANOVA demonstrated significant shifts in relative abundance related to an expansion in *Kazachstania* in HF-fed pigs, but this appears to be somewhat blunted by exposure to LOPs (**Figure 6b**). More precisely, mixed effect regression identified *Kazachstania*, *Candida*, *Cryptococcus*, *Cladosporium*, *Mortierella*, *Aspergillus* and *Penicillium* among the most differentially abundant genera (all FDR-adjusted *p* < 1.0 x10^-11^, **Supplement Table 6**). ANCOM identified significant blooms in *Candida, Kazachstania* and *Torulaspora* on a HF diet (**Supplement Table 7**).

**Figure 6.**
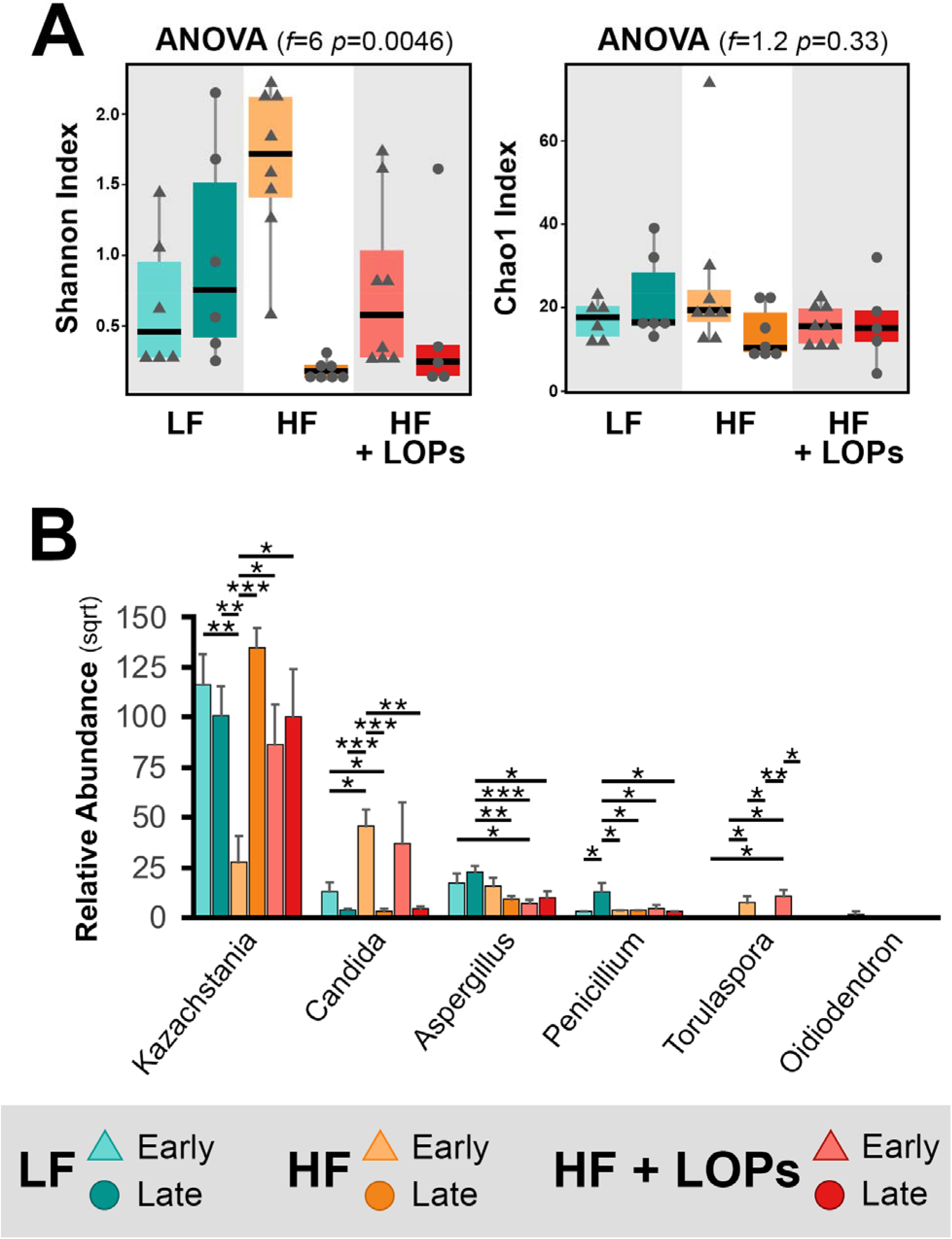
Fungal gut ecology is dominated by a single species and diversity contracts on high fat diet. **a** Fungal diversity as quantified by the Shannon diversity index declines with high fat diet (ANOVA *f* = 6 *p* = 0.0046), but not as quantified by Chao1 (ANOVA *f* = 1.2, *p* = 0.33). **b** Relative abundance of fungal genera alter with time (ANOVA false-discovery rate adjusted \**p* <0.05, \*\**p*<0.01, \*\*\**p* <0.001).

By the final timepoint (after ~5 months on the diet regimen), the mycobiome of LF-fed pigs had diverged slightly from the two groups of high fat fed pigs. Redundancy analysis displays unique but not significantly differential clustering of communities in LF-fed pigs (variance = 9.36, *f* = 1.16, *p* = 0.091) that is not apparent with other techniques (Bray-Curtis dissimilarity, PERMANOVA, *R^2^* = 0.127, *p* = 0.341, PERMDISP2, *p* = 0.28). Diversity was also similar between groups (Shannon, ANOVA, *f* = 3.5, *p* = 0.056; Chao1, ANOVA, *f*= 1.4, *p* = 0.26, **Figure 7a**). Core mycobiome analysis, however, identified two unique genera in animals exposed to LOPs aside from a core of 11 genera (**Figure 7b**). Univariate analysis with ANOVA suggests that differences may be primarily driven by increased abundance of relatively low abundance genera *Wallemia, Penicillium* and *Aspergillus* (**Figure 7c**). Mixed effect regression identified *Kazachstania, Cryptococcus, Aspergillus, Cladosporium, Mortierella, Penicillium, Fusarium* and *Wallemia* as the most significantly differentially abundant genera (FDR-adjusted *p* < 0.00005, **Supplement Table 8**). ANCOM, however, did not identify any differentially abundant genera.

**Figure 7.**
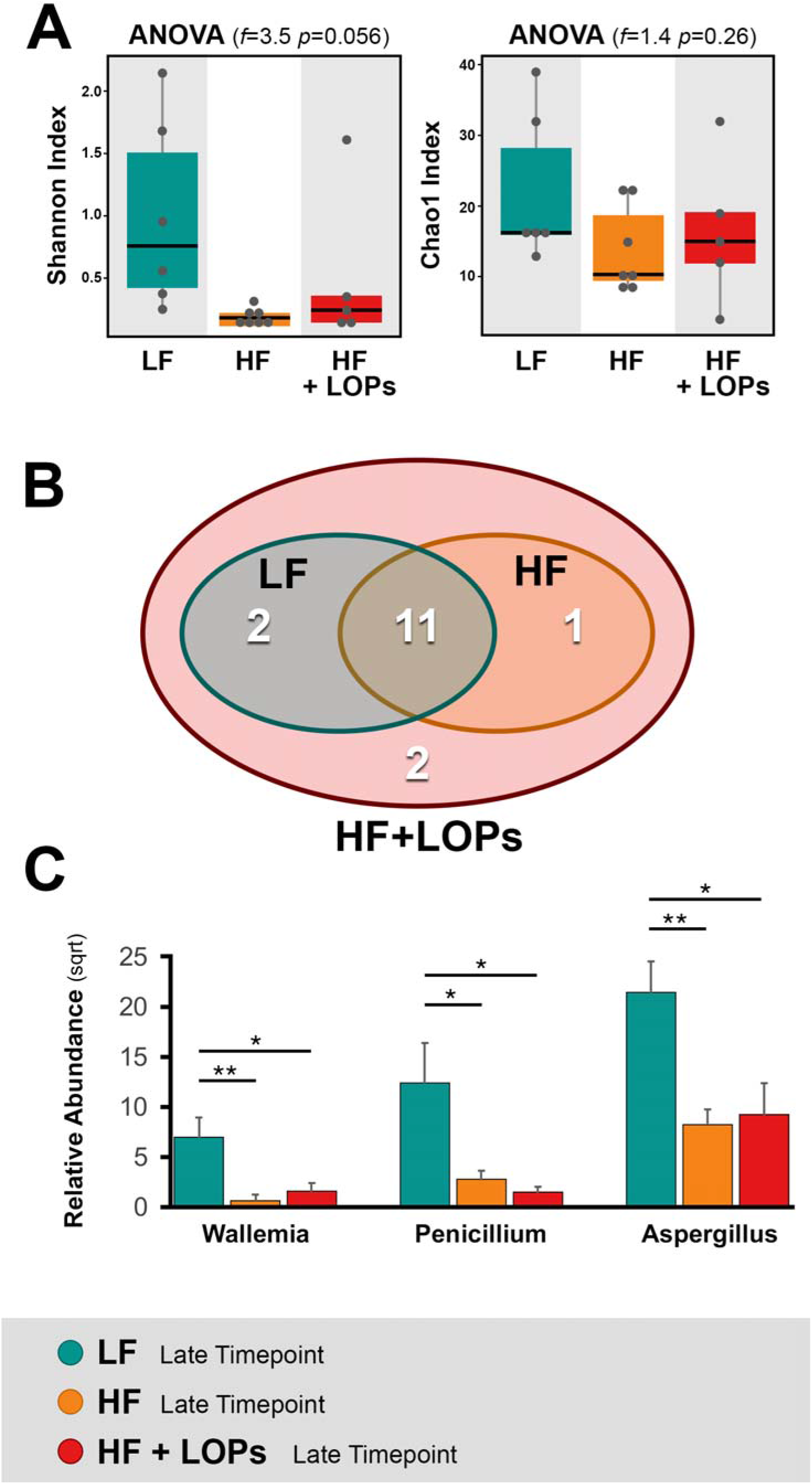
Fungal ecology remains relatively unchanged by dietary interventions. **a** Fungal diversity is also unchanged (Shannon, ANOVA *f* = 3.5, *p* = 0.056; Chao1, ANOVA *f* = 1.4, *p* = 0.26). **b** Core microbiome analysis demonstrates unique fungal genera after lipid oxidation product exposure. **c** Relative abundance of fungal genera (ANOVA false-discovery adjusted \**p* <0.05, \*\**p*<0.01, \*\*\**p* <0.001).

### 3.4 Machine Learning Based Modeling

Outlier detection was performed on the full dataset before model development. Due to limited sample size, three pigs exceeded the outlier threshold for the three dependent variables of interest (TBF, EBF and TrBF). As a result, the full dataset was split into two additional subsets that had outliers removed to minimize outsized influence.

After implementing LOOCV, 18 different scenarios were generated across two taxonomic ranks and three different datasets for the three dependent variables of interests (**Table 2**). For the TBF variable, the use of 19 pigs as opposed to the full dataset yielded the optimal models at both family and genus levels. At the family level, the mean and median of the MSE were reported to be 0.0563 and 0.023, respectively. Similarly, at the genus level, the mean and median of the MSE were reported at 0.0559 and 0.0241, respectively. The MSEs for TBF were greatly reduced after outlier removal as the mean and median of the MSE in the full dataset were 0.4124 and 0.0205, respectively. Similar trends were observed in EF at the family level. Here, the optimal mean and median MSE for EF were 0.0597 and 0.0079, respectively. However, it appears that the dataset with the lowest mean and median MSE at the genus level only required the removal of two outliers as opposed to the three outliers removed in MSE determination for TBF. Mean and median MSE calculations for TRF followed patterns established in TBF MSE calculations. Optimal models generated at the family and genus level emerged from the usage of the smallest dataset. As expected, the mean was largely affected by the inclusion of outliers. In the full dataset, the mean and median MSE for TRF was 0.7179 and 0.035, respectively at the family level. This contrasts to the smallest dataset where the mean and median MSE were reported to be 0.0887 and 0.0398, respectively at the genus level. Outlier detection highlights the outsized influence of outliers on smaller datasets. There was more congruency between true and predicted values for each dependent variable once the outliers were removed from the full dataset.

**Table 2.**
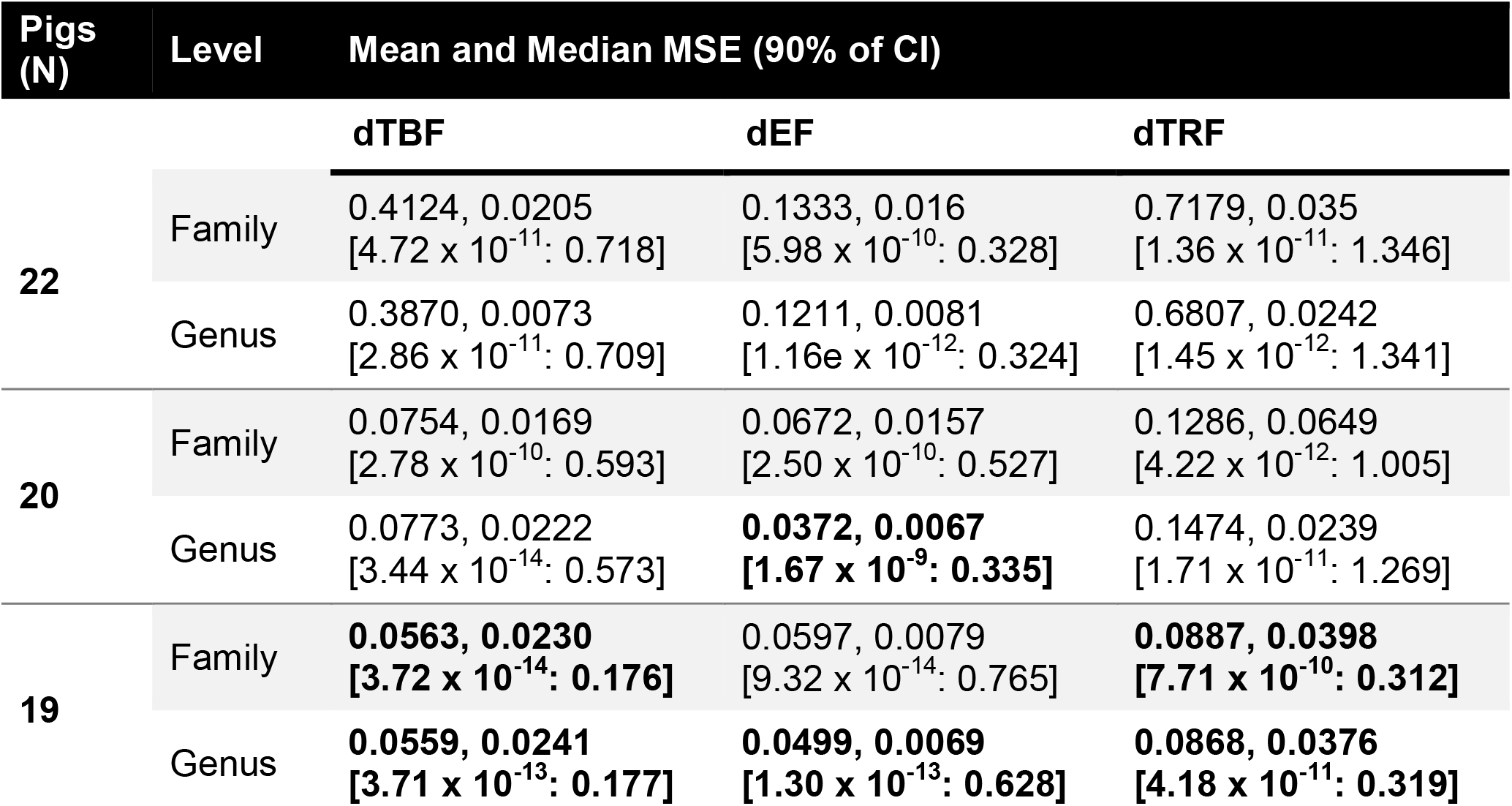
Mean and Median of MSE (90% of CI) scores for 18 different LOOCV scenarios. Calculations of mean squared error (MSE) of test sample of each fold. Mean and median of MSE of each LOOCV fold along with 90% confidence intervals (CI) based on quarters. Bolded values represent the optimal predictive model combinations for each dependent variable.

| Pigs<br>(N) | Level | Mean and Median MSE (90% of CI) |  |  |
| --- | --- | --- | --- | --- |
|  |  | dTBF | dEF | dTRF |
| 22 | Family | 0.4124, 0.0205<br>[4.72 x 10<sup>-11</sup>: 0.718] | 0.1333, 0.016<br>[5.98 x 10<sup>-10</sup>: 0.328] | 0.7179, 0.035<br>[1.36 x 10<sup>-11</sup>: 1.346] |
|  | Genus | 0.3870, 0.0073<br>[2.86 x 10<sup>-11</sup>: 0.709] | 0.1211, 0.0081<br>[1.16e x 10<sup>-12</sup>: 0.324] | 0.6807, 0.0242<br>[1.45 x 10<sup>-12</sup>: 1.341] |
| 20 | Family | 0.0754, 0.0169<br>[2.78 x 10<sup>-10</sup>: 0.593] | 0.0672, 0.0157<br>[2.50 x 10<sup>-10</sup>: 0.527] | 0.1286, 0.0649<br>[4.22 x 10<sup>-12</sup>: 1.005] |
|  | Genus | 0.0773, 0.0222<br>[3.44 x 10<sup>-14</sup>: 0.573] | **0.0372, 0.0067**<br>**[1.67 x 10<sup>-9</sup>: 0.335]** | 0.1474, 0.0239<br>[1.71 x 10<sup>-11</sup>: 1.269] |
| 19 | Family | **0.0563, 0.0230**<br>**[3.72 x 10<sup>-14</sup>: 0.176]** | 0.0597, 0.0079<br>[9.32 x 10<sup>-14</sup>: 0.765] | **0.0887, 0.0398**<br>**[7.71 x 10<sup>-10</sup>: 0.312]** |
|  | Genus | **0.0559, 0.0241**<br>**[3.71 x 10<sup>-13</sup>: 0.177]** | **0.0499, 0.0069**<br>**[1.30 x 10<sup>-13</sup>: 0.628]** | **0.0868, 0.0376**<br>**[4.18 x 10<sup>-11</sup>: 0.319]** |

Variable importance analysis revealed consistent patterns across all three dependent variables. As expected, dietary regimen, particularly variability between LF, HF, HF+LOPs, were identified as important variables in explaining variation between the dependent variables. Additionally, the sex independent variable was determined to be important in predicting the variation in the dependent variables. Interestingly, the microbial species belonging to the *Methanobacteria* order (*Methanobacteriaceae, methanomassiliicoccaceae*, and *methanosphera*) were the only microorganisms that were consistently identified to be important predictors of variability in body fat (TBF, EF and TrF; Importance > 20%; **Figure 8**).

**Figure 8.**
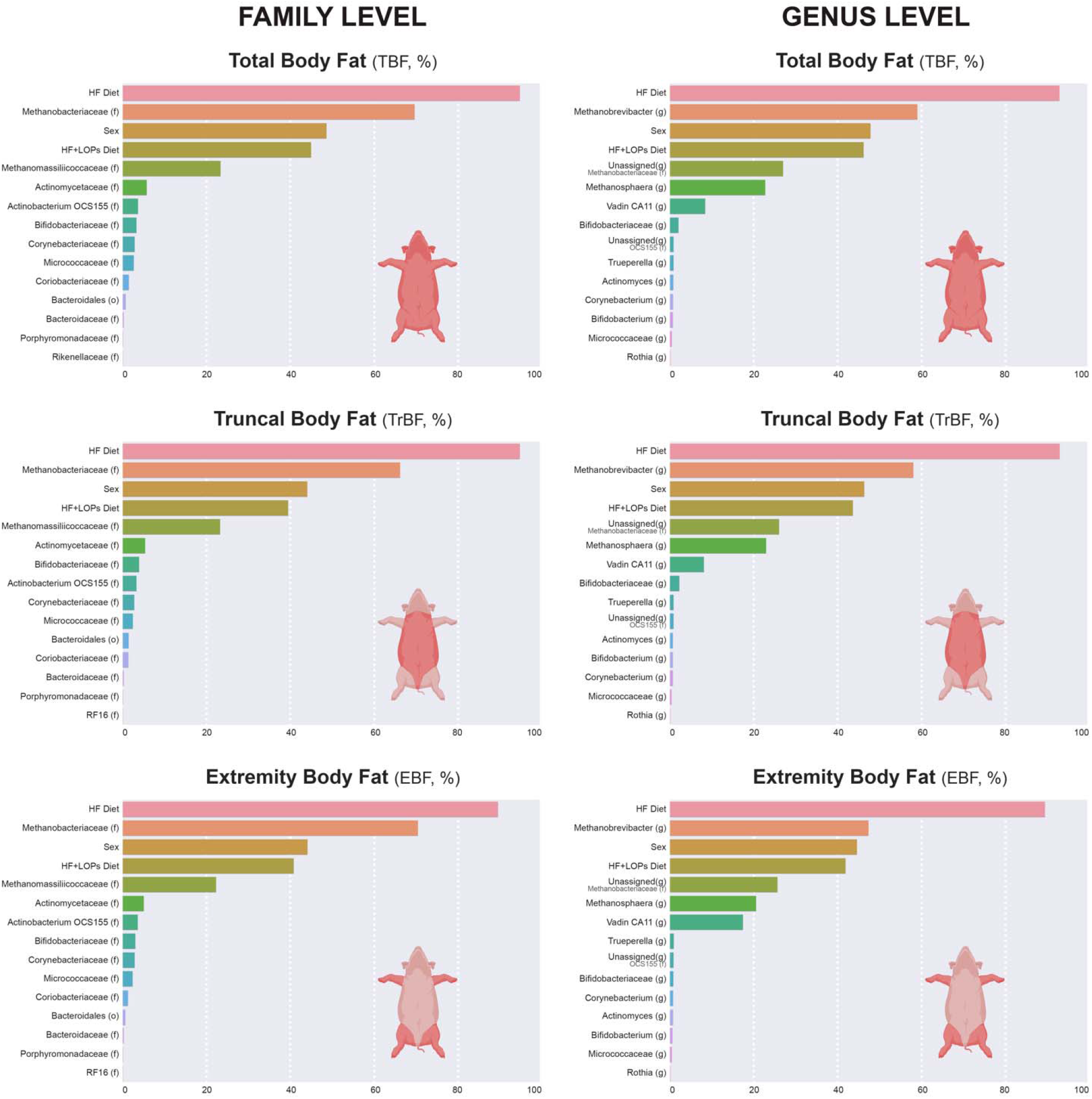
Variable importance analysis on final Light Gradient Boosting based models. The importance of variables calculated based on the “gain” importance variable importance analysis of the LightGBM model. The bars associated with the variables represents the relative contribution of the variables to the prediction, where longer bars imply higher importance.

## 4. DISCUSSION

Despite the ubiquitous nature of LOPs in the Westernized diet, little is known about the effects of consuming these compounds on human biology. Increased fried food consumption is not limited to the United States and is observed in other developed and developing countries.^52–57^ These diets are increasingly consumed by children and adolescents, ^17,23,58,59^ raising concerns regarding how factors that influence chronic disease may be affected. In this study, we utilized a pig model to examine the effects of LOPs in high-fat diets on growth, body composition and gut microbiome during the pre-pubescent period. The major findings of the study are that the consumption of LOPs produces (1) decreased growth (weight gain) and adiposity and (2) dysbiosis of the gut microbiome, with a strong association between the two.

The consumption of high fat diets by pigs in the prepubertal period led to higher body weights and % total body fat, when compared to those consuming a low-fat (LF) diet. However, the presence of LOPs in the high fat diet (HF+LOPs), led to lower growth rates and adiposity compared to a high fat diet with low amounts of LOPs (HF). Pigs used in our study were obtained from our breeding and maintenance facility that houses approximately 1,500 pigs at a given time. Here, the pigs reach sexual maturity at approximately 5.5 months of age, which corresponds to the end of our study period. This is the first animal study in which the effects of LOPs on growth and adiposity were explored in the context of translational relevance to humans. Previous research studies of LOPs in pigs in the field of livestock agriculture do exist and allow us to draw some comparisons to our findings.

Fats and oils are added to feed in swine livestock production to provide energy for optimal growth.^60^ However, in the United States, in order to increase profit margins, low-cost sources of fats and oils such as spent restaurant oils and dried distillers grains with solubles (DDGS, “waste stream” of various other industries are used).^61–63^ DDGS and spent restaurant oils can introduce large quantities of LOPs into livestock (e.g., swine) production diets.^64,65^ The impact of such widely-used sources of fats on efficiency of swine production and the health of the animals have been extensively studied.^66^ Many of these studies have observed that feeding of LOPs can significantly diminish animal growth.^67–72^ The findings of our study are generally consistent with these studies in the livestock agriculture field. However, the effects of LOPs on body weight in our study are modest compared to several above-mentioned studies. For example, Lu and colleagues found that pigs fed diet containing 5% oxidized oil resulted in a 39.4% reduction in body weight compared to those fed a diet containing unoxidized oil.^72^ Collectively, these observations suggest the oxidative state of dietary lipids may have important implications for childhood growth and development.

The mechanisms by which exogenous sources of LOPs impact growth and body fat have not been well studied. Given that these compounds in the diet are likely to be present at relatively higher concentrations in the gastrointestinal tract than in circulation, any observed gastrointestinal effect may, at least in part, be responsible for a systemic effect. Dibner and colleagues observed that pigs fed LOPs had increased enterocyte turnover rates, proliferation and migration to the gastrointestinal lumen that is indicative of a reduced enterocyte half-life and increased gastrointestinal damage.^66^ In our study, we found secondary LOPs such as acrolein, hexanal, 2-hexanal, 2,4-decadienal, pentanal, 2-heptenal and 4-HNE to be present in notably higher concentrations in our spent restaurant oil when compared to the unoxidized corn oil (**Table 2**). Hazuka and colleagues showed that oil extracted from potato fritters deep fried in cooking oil that was used over 1-5 day of frying, as often done in restaurants, contained PV of 10.3 - 11.5 mEq O_2_/kg of extracted oil and AV of 103 - 184.^73^ Thus, the oil used to provide LOPs in our study is in the ballpark for representing LOPs level from human fast-food item such as potato fries or fritters, although the PV (28.68 mEq O_2_/kg) was higher while AV (45.13) was lower in our LOPs source. Another study found that oil extracted from fried potatoes contained approximately 536 mg/L of HNE in the oil,^74^ in contrast to the lower 15.0 mg/L of HNE in our LOPs source. A number of *in vitro and in vivo* studies provide evidence that specific LOPs can exert a wide range of hazardous effects.^75–82^ A study examining the effects of acrolein on the gut in mice observed that the compound disrupts tight junction proteins and causes epithelial cell death leading to intestinal barrier dysfunction and permeability.^81^ Given that increases in oxidative stress in the gut can impair gut function and nutrient absorption,^83^ we questioned whether LOPs would influence the gut microbiota and whether the effects on the microbiome were associated with the changes in body composition, especially given the current interest in the link between adiposity and gut microbiome.^84–87^

The gut hosts an ecologically rich microbial community comprising of several domains. Bacterial communities in the gut have been studied extensively over the past decade and have been found to have a pivotal role in host immune defense, metabolism and health. The mycobiome (fungal communities), on the other hand, remains largely unstudied although it has gained attention in the last decade.^88,89^ The composition of the microbiota is dynamic and can be modulated by internal and external stimuli such as local inflammatory state, environment, host genetics and diets.^90–93^ The disruption of the microbiome community structure and function, referred to as dysbiosis, is observed in many diseases such as inflammatory bowel disease, obesity and type 2 diabetes.^30,94,95^ Although, there is ample evidence that dietary fat intake causes dysbiosis of the gut bacteriome and mycobiome,^30,93,96^ no study to date has given any attention to the oxidative state of the fat.

We report here, for the first time, that dietary LOPs uniquely impact gut microbial ecology and explore the association between these alterations and body composition. Bacteriome and mycobiome were examined at 2 months (early timepoint) and approximately five months (late timepoint) after beginning dietary regimens. Over the course of all dietary regimens, shifts in bacterial alpha and beta diversity were observed (**Figure 4**). An age-related decline in diversity has been previously reported in humans, porcine and rodent studies, with a greater decrease seen in subjects consuming a high-fat diet.^97–100^ A lower gut diversity is widely associated with obesity, metabolic syndrome, diabetes and other chronic diseases.^101–105^ In the current study, dietary fat intake also produced a greater decline in diversity, when compared to the LF diet. The magnitude of the effect was greater in HF+LOPs pigs but did not significantly differentiate from changes in HF pigs (**Figures 4 and 5**).

Levels of *Spirochaetales, RF39, Erysipelotrichales*, and *Lactobacillales* inversely correlated with the HF diets (at both timepoints), regardless of LOP content. These results are consistent with two other porcine studies that demonstrated relative increases in the *Lactobacillales* order was associated with a leaner phenotype exhibited by pigs receiving a low-fat diet.^31,106^ A distinct microbial signature was observed in pigs consuming a high fat diet containing LOPs, including increased *Barnesiellaceae spp*., which is associated with Westernized diet containing limited fiber and in humans with sedentary lifestyles. ^107,108^ *Acinetobacter* and *Rheinheimera* genera were also significantly enriched under dietary LOPs. In mice, the genera *Acinetobacter* is considered to be conditionally pathogenic and is associated with high fat diet-induced microbiota dysbiosis.^109^

There is little previous characterization of the mycobiome in the porcine gut.^44^ Our mycobiome analyses revealed lower community diversity across all dietary regimens than bacterial communities. Prior porcine studies show the mycobiome is dominated by the yeast *Kazachstania*, consistent with our findings.^44,51,110^ Little is known about the symbiotic relationship between the *Kazachstania* genus and diverse bacterial communities or about its role in modulating growth, metabolism, and immune function in pigs. Similar to bacteriome analyses, over the course of the experiment, distinct clustering is observed based on dietary fat intake and levels of LOPs. Significant blooms in *Candida, Kazachstania* and *Torulaspora* were identified in the high fat diet groups. Similarly, *Heisel et al.* reported that the *Candida albicans* spp. exhibited a 16.5% higher abundance in mice fed a high fat chow compared to standard chow.^111^ The dynamic interactions between fungi, bacteria and the host altered by high dietary fat intake remain an area of active exploration.

We developed a gradient boosting machine learning prediction model, determined to be 95% accurate, to identify relevant factors modulating total and regional body fat. Expectedly, dietary fat intake was found to be the most important predictor of body fat (**Figure 8**). Interestingly, the presence of dietary LOP intake was also recognized as an important predictor. However, contrary to what we hypothesized – that increasing LOPs in the western diets may be responsible for increasing incidence of obesity – the presence of LOP in the high fat diet produced a reduction in body fat. While this was unexpected, it is consistent with the finding of *Penumetcha* and colleagues,^112^ in which the authors analyzed human subjects data from two cycles of National Health and Nutrition Examination Surveys, with a survey sample of 9,982 subjects between 2 and 85 years of age and observed inverse correlation between LOPs intake and obesity but positive associations with glucose intolerance. Their observation is further supported by a study in rats and mice that found that the consumption of oxidized frying oil (prepared by frying wheat flour dough sheets in soybean oil heated to ~205°C in the laboratory) is less adipogenic, but induces glucose intolerance.^113^ These observations contribute to the concept of metabolically healthy obesity, where adiposity alone is not sufficient or necessary for glucose intolerance or insulin resistance.

The variable importance variable analysis on the final machine learning models also identified the abundance of methanogens, specifically *Methanobacteriaceae* and *Methanomassiliicoccaceae* families, as predictors of total and region-specific body fat (**Figure 8**). The genus *Methanobrevibacter* was identified as one of the strongest predictors, having an importance level second to the fat content of the diet. While *Methanobrevibacter* is recognized as major methanogen in the gut of pigs,^114,115^ little is known about its influence over adiposity in pigs.^116^ However, in prepubescent humans, Methanocrevibacter colonization was associated higher weight z-scores, BMI, and adiposity.^117^ In adult patients, *Zhang et al*. observed that obese individuals have significantly higher abundances of members of the order *Methanobacteriales* than post-gastric bypass and normal individuals.^118^

LOPs have become prevalent in human^119^, livestock ^67–72^ and pet^120^ diets in the United States and other developed countries. With the exception of research in livestock agriculture field focusing on growth and meat quality, there is little research on the impact of dietary LOPs, especially as it relates to human or other animal species. A study examining the effects of LOPs in pet food, showed that presence of LOPs reduced total body fat content, bone development and immune function in growing puppies, suggesting the relevance of dietary LOPs to companion animal health.^121^ Our study demonstrates that while unoxidized dietary fat intake during the pre-pubertal period is associated with greater growth rate compared to low fat diets, the presence of LOPs notably oppose these effects. High dietary fat intake distinctly altered gut microbiome during this period, with the presence of LOPs producing unique differences separate from those produced by the high fat diet. Machine learning based modeling revealed *Methanobrevibacter* to be a very important predictor of body fat. Further studies in translational animal models such as porcine models, are needed to not only understand this link, but also to explore the broader implication of dietary LOPs on human biology, especially during the early developmental period, given the still increasing consumption of these compounds.

## Supporting information

Supplemental Data Index

## ACKNOWLEDGEMENTS

This work was supported by the National Institutes of Health [NCI, R01 CA253329; NHLBI: K08 HL151907]; National Institute of Food and Agriculture, United States Department of Agriculture [NIFA (Formula Project Grant), WIS01972], and University of Wisconsin-Madison [Graduate Research Scholars Fellowship (Science and Medicine Graduate Research Scholars); Biomedical & Genomic Research Group Discretionary Fund].

We would like to thank Ms. Jennifer Meudt for her technical contributions and her logistical guidance. We also would also like to thank the staff (Mr. Jamie Reichert, Dr. Ana Cecilia Escobar López, Mr. Sam Trace and Ms. Keri Graff) of the Swine Research and Teaching Center (SRTC), the staff (Catherine T. Jobsis and John Kemper) of the Translational Research Facility, and the Attending Veterinarians (Dr. Michael Maroney and Dr. Kay Nelson) at the University of Wisconsin-Madison for their overall programmatic support of the current research. Additionally, we would like to thank Mr. Peter Crump and Mr. Nicholas Keuler of the Department of Statistics for consultation on the statistical analyses. Images of pigs used in figures were created with Biorender.com.

## AUTHOR CONTRIBUTIONS

F.K. Arowolo and D. Shanmuganayagam conceptualized and designed the study with substantial contributions from I. Karabayir (machine-learning), O. Akbiligic (machine-learning), J. Booth (diet design/formulation), and J.F. Pierre (microbiome); F.K. Arowolo, M. Blaser, and J. Booth performed research; F.K. Arowolo, K.A. Willis, I. Karabayir, Oguz Akbiligic, M. Blaser, J.F. Pierre and D. Shanmuganayagam analyzed and interpreted data; F.K. Arowolo, K.A. J.F. Pierre and D. Shanmuganayagam wrote the paper with Willis, I. Karabayir, Oguz Akbiligic, M. Blaser contributing critical revisions. All authors approved the final version.

## ABBREVIATIONS

ANCOM: analysis of comparison of microbiomes
ANOVA: analysis of variance
AV: p-anisidine value
DXA: dual energy x-ray absorptiometry
EBF: extremity body fat
EBM: extremity bone mass
ELM: extremity lean mass
HF: high fat diet with negligible/low lipid oxidation products
HF+LOPs: high fat with high lipid oxidation products diet
LC-MS: liquid chromatography-mass spectrometry
LF: low-fat diet
LightGBM: Light Gradient Boosting Machine
LOOCV: Leave One Out Cross Validation
LOPs: lipid oxidation products
LSM: least-square means
MSE: mean squared error
PCA: Principal components analysis
PCoA: principal coordinates analysis
PERMANOVA: permutational multivariate analysis of variance
PERMDISP2: permutational analysis of multivariate dispersions
PV: peroxide value
RDA: redundancy analysis
TBF: total body fat
TBM: total bone mass
TLM: total lean mass
TrBF: truncal body fat
TrBM: truncal bone mass
TrLM: truncal lean mass

## Supplementary Files

**Supplementary Table 1.**
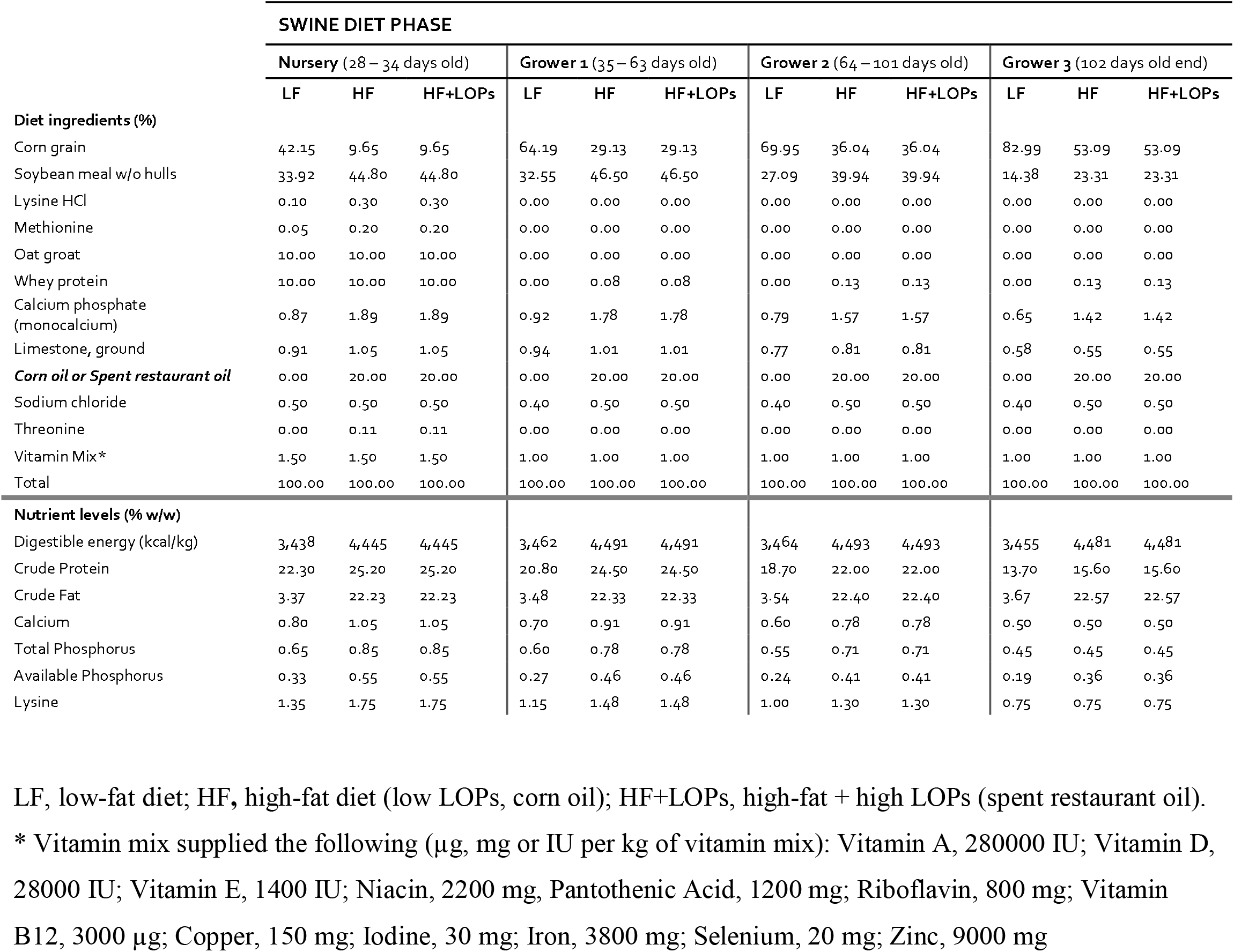
Dietary composition of dietary regimens.

**Supplementary Figure 1.**
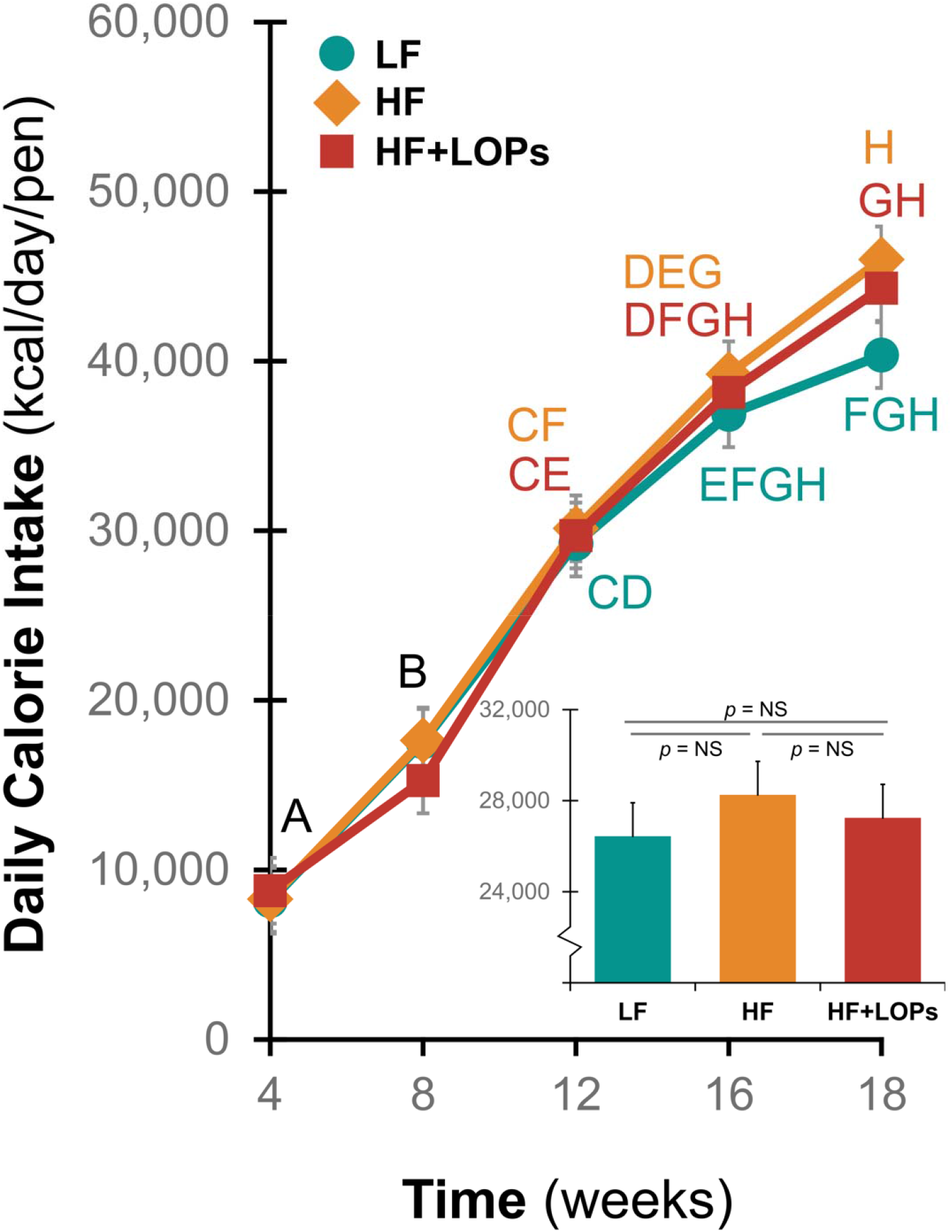
Effects of Dietary Lipid Oxidation Products on Body Weight and Adiposity (Total and Regional) in the Prepubertal Period. The line graphs show actual calorie intake based on feed intake by the dietary groups. Data is presented as least square means ± SEM. Means without any common letter differ, *p* < 0.05. The inset bar graphs show the overall calorie intake (least square means ± SEM). No significant difference in calorie intake was observed between the groups.

**Supplementary Figure 2 and Supplementary Tables 2 - 8 are provided as separate individual files.**

