## Supplemental Data Index for "Dietary Lipid Oxidization Products Alter Growth, Adiposity and Gut Microbial Ecology in Prepubertal Porcine Model"

**Supplemental Data 1.** Dietary composition of dietary regimens.

**Supplemental Data 2.** Daily Calorie Intake Based on Feed Consumption.

**Supplemental Data 3.** Mixed effect regression analysis of 16S taxa across timepoints.

**Supplemental Data 4.** ANCOM analysis of select 16S taxa

**Supplementary Data 5.** Linear Discriminant Analysis of Effect Size (LefSe) was used to identify key taxa that were enriched under experimental diets.

**Supplemental Data 6.** Mixed effect regression analysis of 16S taxa between Week 12 and Week 19.5

**Supplemental Data 7.** ANCOM of *Epulopiscium* at experimental endpoint (Week 19.5).

**Supplemental Data 8.** Mixed effect regression analysis of ITS taxa across timepoints.

**Supplemental Data 9.** ANCOM analysis of select ITS taxa.

**Supplemental Data 10.** Mixed effect regression analysis of ITS taxa between Week 12 and Week 19.5
