## Supplemental Data 1 for "Dietary Lipid Oxidization Products Alter Growth, Adiposity and Gut Microbial Ecology in Prepubertal Porcine Model"

**Supplemental Data 1. Dietary composition of dietary regimens.**

|  | **SWINE DIET PHASE** | | | | | | | | | | | |
| --- | --- | --- | --- | --- | --- | --- | --- | --- | --- | --- | --- | --- |
|  | **Nursery** (28 – 34 days old) | | | **Grower 1** (35 – 63 days old) | | | **Grower 2** (64 – 101 days old) | | | **Grower 3** (102 days old end) | | |
|  | **LF** | **HF** | **HF+LOPs** | **LF** | **HF** | **HF+LOPs** | **LF** | **HF** | **HF+LOPs** | **LF** | **HF** | **HF+LOPs** |
| **Diet ingredients (%)** |  |  |  |  |  |  |  |  |  |  |  |  |
| Corn grain | 42.15 | 9.65 | 9.65 | 64.19 | 29.13 | 29.13 | 69.95 | 36.04 | 36.04 | 82.99 | 53.09 | 53.09 |
| Soybean meal w/o hulls | 33.92 | 44.80 | 44.80 | 32.55 | 46.50 | 46.50 | 27.09 | 39.94 | 39.94 | 14.38 | 23.31 | 23.31 |
| Lysine HCl | 0.10 | 0.30 | 0.30 | 0.00 | 0.00 | 0.00 | 0.00 | 0.00 | 0.00 | 0.00 | 0.00 | 0.00 |
| Methionine | 0.05 | 0.20 | 0.20 | 0.00 | 0.00 | 0.00 | 0.00 | 0.00 | 0.00 | 0.00 | 0.00 | 0.00 |
| Oat groat | 10.00 | 10.00 | 10.00 | 0.00 | 0.00 | 0.00 | 0.00 | 0.00 | 0.00 | 0.00 | 0.00 | 0.00 |
| Whey protein | 10.00 | 10.00 | 10.00 | 0.00 | 0.08 | 0.08 | 0.00 | 0.13 | 0.13 | 0.00 | 0.13 | 0.13 |
| Calcium phosphate (monocalcium) | 0.87 | 1.89 | 1.89 | 0.92 | 1.78 | 1.78 | 0.79 | 1.57 | 1.57 | 0.65 | 1.42 | 1.42 |
| Limestone, ground | 0.91 | 1.05 | 1.05 | 0.94 | 1.01 | 1.01 | 0.77 | 0.81 | 0.81 | 0.58 | 0.55 | 0.55 |
| ***Corn oil or Spent restaurant oil*** | 0.00 | 20.00 | 20.00 | 0.00 | 20.00 | 20.00 | 0.00 | 20.00 | 20.00 | 0.00 | 20.00 | 20.00 |
| Sodium chloride | 0.50 | 0.50 | 0.50 | 0.40 | 0.50 | 0.50 | 0.40 | 0.50 | 0.50 | 0.40 | 0.50 | 0.50 |
| Threonine | 0.00 | 0.11 | 0.11 | 0.00 | 0.00 | 0.00 | 0.00 | 0.00 | 0.00 | 0.00 | 0.00 | 0.00 |
| Vitamin Mix* | 1.50 | 1.50 | 1.50 | 1.00 | 1.00 | 1.00 | 1.00 | 1.00 | 1.00 | 1.00 | 1.00 | 1.00 |
| Total | 100.00 | 100.00 | 100.00 | 100.00 | 100.00 | 100.00 | 100.00 | 100.00 | 100.00 | 100.00 | 100.00 | 100.00 |
| **Nutrient levels (% w/w)** |  |  |  |  |  |  |  |  |  |  |  |  |
| Digestible energy (kcal/kg) | 3,438 | 4,445 | 4,445 | 3,462 | 4,491 | 4,491 | 3,464 | 4,493 | 4,493 | 3,455 | 4,481 | 4,481 |
| Crude Protein | 22.30 | 25.20 | 25.20 | 20.80 | 24.50 | 24.50 | 18.70 | 22.00 | 22.00 | 13.70 | 15.60 | 15.60 |
| Crude Fat | 3.37 | 22.23 | 22.23 | 3.48 | 22.33 | 22.33 | 3.54 | 22.40 | 22.40 | 3.67 | 22.57 | 22.57 |
| Calcium | 0.80 | 1.05 | 1.05 | 0.70 | 0.91 | 0.91 | 0.60 | 0.78 | 0.78 | 0.50 | 0.50 | 0.50 |
| Total Phosphorus | 0.65 | 0.85 | 0.85 | 0.60 | 0.78 | 0.78 | 0.55 | 0.71 | 0.71 | 0.45 | 0.45 | 0.45 |
| Available Phosphorus | 0.33 | 0.55 | 0.55 | 0.27 | 0.46 | 0.46 | 0.24 | 0.41 | 0.41 | 0.19 | 0.36 | 0.36 |
| Lysine | 1.35 | 1.75 | 1.75 | 1.15 | 1.48 | 1.48 | 1.00 | 1.30 | 1.30 | 0.75 | 0.75 | 0.75 |

LF, low-fat diet; HF**,** high-fat diet (low LOPs, corn oil); HF+LOPs, high-fat + high LOPs (spent restaurant oil).

* Vitamin mix supplied the following (µg, mg or IU per kg of vitamin mix): Vitamin A, 280000 IU; Vitamin D, 28000 IU; Vitamin E, 1400 IU; Niacin, 2200 mg, Pantothenic Acid, 1200 mg; Riboflavin, 800 mg; Vitamin B12, 3000 µg; Copper, 150 mg; Iodine, 30 mg; Iron, 3800 mg; Selenium, 20 mg; Zinc, 9000 mg
