## Supplemental Data 2 for "Dietary Lipid Oxidization Products Alter Growth, Adiposity and Gut Microbial Ecology in Prepubertal Porcine Model"

**Supplemental Data 2. Daily Calorie Intake Based on Feed Consumption.**

The line graphs show actual calorie intake based on feed intake by the dietary groups. Data is presented as least square means ± SEM. Means without any common letter differ, *p* < 0.05. The inset bar graphs show the overall calorie intake (least square means ± SEM). No significant difference in calorie intake was observed between the groups.

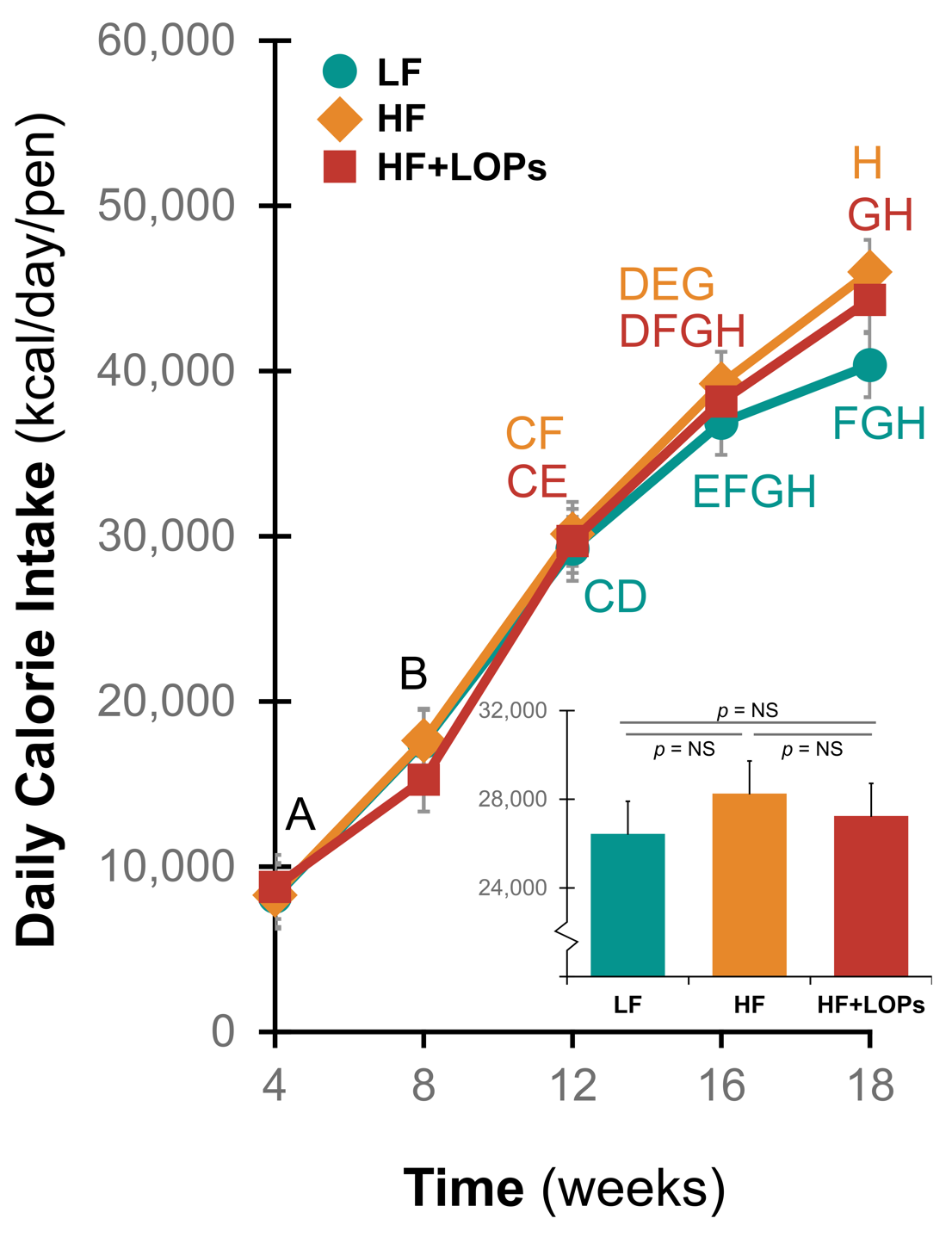
