## Supplemental Data 5 for "Dietary Lipid Oxidization Products Alter Growth, Adiposity and Gut Microbial Ecology in Prepubertal Porcine Model"

**Supplementary Data 5. Linear Discriminant Analysis of Effect Size (LefSe) was used to identify key taxa that were enriched under experimental diets.**

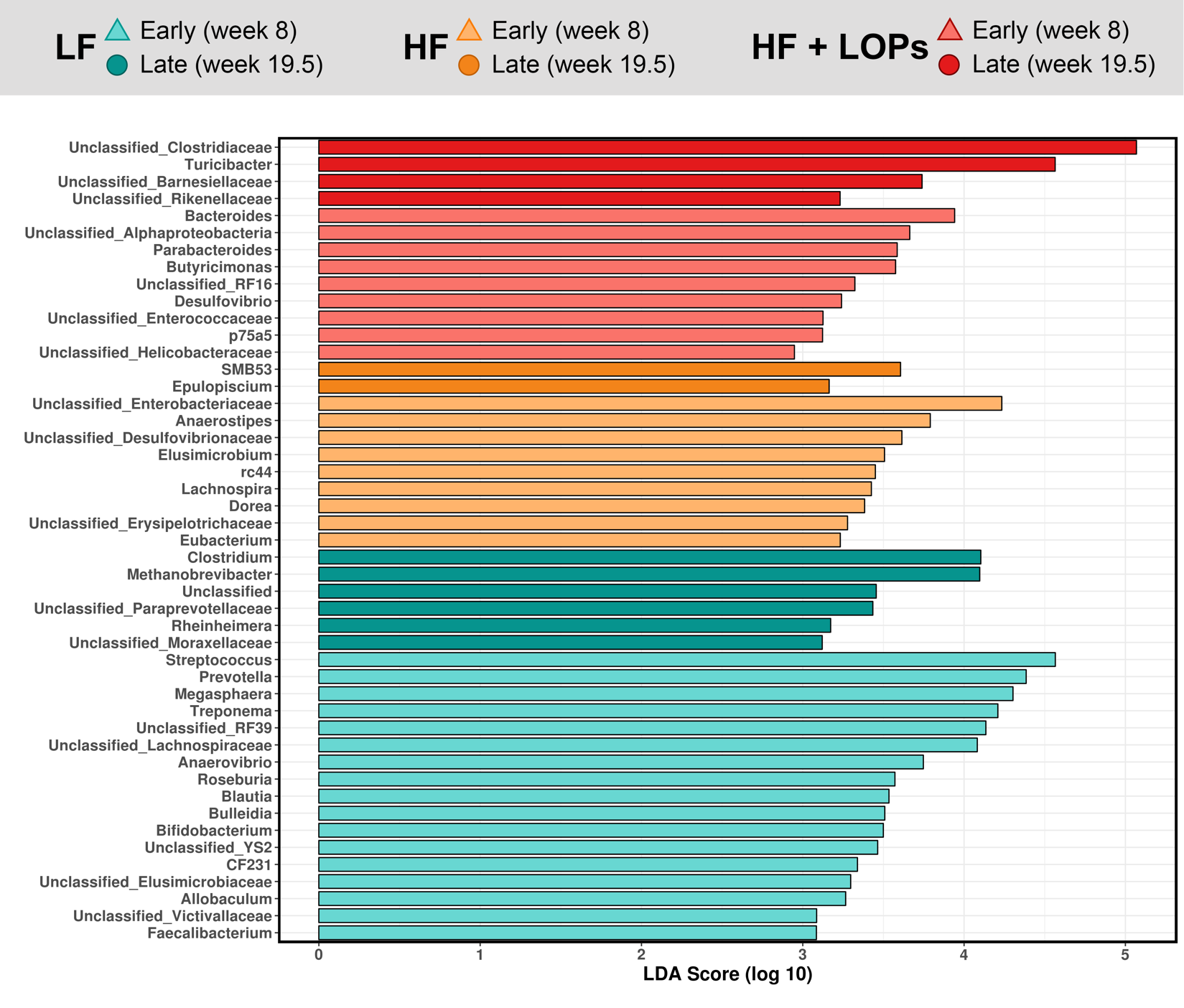
